# PanScan: A tertiary analysis tool for pangenome graph

**DOI:** 10.1101/2025.05.01.651685

**Authors:** Bipin Balan, Shehzad Hanif, Mohammad Amiruddin Hashmi, Muhammad Kumail, Shuhd BinEshaq, Talal Al-Yazeedi, Richa Tambi, Munazza Murtaza, Bassam Jamalalail, Hanan Elsokary, Maryam Al Obathani, Nesrin Mohamed, K Dasuki Hansana Perera, Dia Advani, Aysha Ibrahim Hamzah, Suhana Shiyas, Nelson C. Soares, Omer AlKhnbashi, Stefan S Du Plessis, Mohamed Almarri, Nasna Nassir, Alawi Alsheikh-Ali, Mohammed Uddin

## Abstract

The genomic representation of populations across the globe is critical to ensuring a comprehensive and equitable human reference. Constructing a pangenome graph reference for different populations is the best approach to addressing local genomic diversities. Although major initiatives across continents are underway to construct pangenome graph references, the field lacks the necessary toolsets for tertiary analysis to characterize telomere-to-telomere (T2T) assemblies and the complexity of haplotypes. PanScan is a bioinformatics software package developed for human pangenome tertiary analysis. It includes multiple modules designed to detect duplicated gene sets from T2T assemblies, identify novel variants and sequences, as well as detect and visualize complex genomic regions through pangenome graph haplotype loops. We have used multiple pangenomes across different populations to assess the tertiary analysis and their accuracy. The tool is designed to streamline tertiary analysis and is compatible with multiple pangenome graph construction algorithms. PanScan is freely available on GitHub (https://github.com/CATG-Github/panscan), where users can provide human pangenome assemblies or VCF files as inputs for automated analyses through command-line operations on Linux systems.

**Graphical Abstract:** 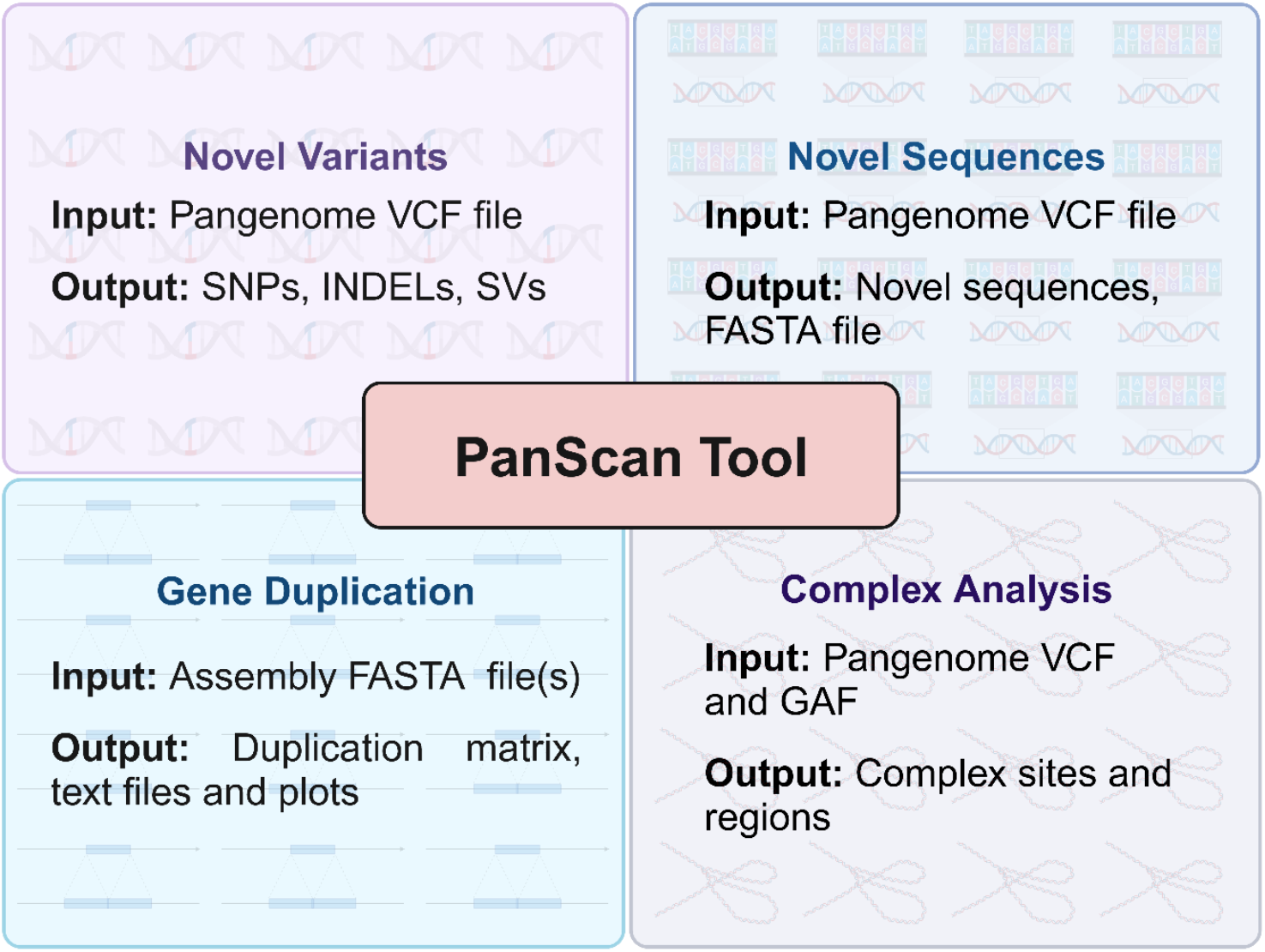

## 1. Introduction

The advent of high-throughput human genome sequencing has revolutionized our understanding of normal biology and disease origins, transforming the human pangenome into a groundbreaking resource that is reshaping biomedical research (1). Unlike traditional linear reference genomes, a pangenome represents a collective genomic architecture that integrates genes, regulatory elements, and non-coding regions across ethnically diverse individuals. Although genes contribute as the foundational basis for essential biological functions, new haplotypes with population specific alleles often reflecting adaptive diversity driven by structural variations (SVs), insertion/deletion (INDELs), single nucleotide polymorphisms (SNPs), and novel sequences. Genomic diversity shaped by demographic history and selective pressures, underscores the necessity of a pangenome framework to resolve the full spectrum of human genetic variation. This approach enables precise allele identification, evolutionary lineage tracking, and trait-associated variant discovery (2–4). The clinical implications of pangenome analysis are substantial and far-reaching. Conserved regions offer stable therapeutic targets for broad-spectrum interventions, whereas variable loci guide personalized medicine strategies tailored to population-specific disease susceptibilities, such as oncogenic structural rearrangements or pharmacogenomic variants (5, 6). Furthermore, pangenomic insights are driving CRISPR-Cas9-based gene-editing technologies, facilitating targeted disease interventions and engineered cell therapies by mapping conserved functional domains and context-dependent regulatory network (7–9). Despite milestones like the Human Pangenome Reference Consortium (HPRC) (10), Chinese Pangenome Consortium (CPC) (11), and Arab Pangenome Reference (APR) (12), a gap persists in resolving population-specific variabilities. This necessitates geographically stratified pangenomes to capture underrepresented alleles and cryptic SVs (10). Advances in long-read sequencing now enable high-contiguity assemblies, making large-scale, population-inclusive pangenome construction feasible while also capturing a more detailed depiction of population-specific traits, rare genetic variants, and previously unidentified gene functions (11, 13). Although the pangenome approach provides valuable insights into genomic diversity, the complexity of graph representation has made clinical adaptation challenging, primarily due to the lack of automated tools that enable the accurate and intuitive interpretation of genomic variations.

Parallel to these developments, the emergence of pangenome graphs has significantly advanced the analysis of complex genomic data. A pangenome graph provides a graphical representation of multiple aligned genomes, efficiently capturing genetic variations such as SVs, INDELs, and SNPs. Complementing this advancement, a diverse suite of computational tools has emerged to facilitate pangenome assembly, alignment, variant analysis, and visualization. Tools such as Anvi’o, PanTools, and panX are particularly useful for analysing the complex structure of human pangenomes and visualizing gene annotation and diversity (14, 15). Other specialized tools address key methodological gaps by performing specific tasks within pangenome analysis. For instance, Pandagma assists in identifying gene duplication events (16), while PanGraph, VG (Variant Graph), and PASA enable the identification of novel sequences, SVs, and transcriptome-level investigations, respectively (17, 18). Likewise, SyRI and mummer4 streamline the detection of complex rearrangements in highly dynamic genomic regions (19, 20). Despite the availability of these various analytical tools, integrating them into a unified pipeline remains a labour-intensive and time-consuming process, often necessitating manual data harmonization, thereby limiting reproducibility. Due to the complexity of tertiary pangenome analysis, along with its dependence on multiple independent tools that each present unique installation and operational challenges, highlights the need for a more streamlined and automated approach.

PanScan is a comprehensive computational tool that offers advanced capabilities for identifying novel sequences, SVs, and repetitive regions from T2T assemblies or pangenome graph. With its ability to annotate duplicate genes and visualize complex genomic landscapes, PanScan provides invaluable insights into genetic diversity. By streamlining analysis and reducing manual effort, it enhances research efficiency and facilitate large-scale genomic studies. Consequently, PanScan serves as a critical tool for advancing human pangenomic research, identifying population-specific genetic variations and will accelerate the adaptation of graph pangenome into clinical setting.

## 2. Method

### 2.1. GIAB dataset: Pangenome construction

To demonstrate the use case and the utility of the multiple built-in modules of the PanScan tool, we used three trios from the Genome in a Bottle (GIAB) dataset (HG002, HG005, and NA19240) which were not included in the HPRC pangenome. Using Minigraph-Cactus (v2.9.3) (21), we integrated six (haplotypes) fully phased long-read (PacBio HiFi) assemblies into a single graph structure. CHM13 was used as the backbone, supplemented by the GRCh38 reference genome and subsequently expanded using Minigraph to incorporate the new assemblies. The methodology for building the pangenome is detailed in our previous work (12).

### 2.2. PanScan Tool

To facilitate the tertiary analysis of graph-based human pangenomes, we developed PanScan, a comprehensive computational tool for identifying and visualizing complex structural regions within pangenome graphs. PanScan enables the detection and annotation of gene duplications from assemblies and identifies novel variants and sequences by comparing them against existing databases. The methodology for each functionality is summarized in the workflow diagram (Figure 1, Table 1), with detailed explanation provided in the subsequent sections.

**Table 1:**
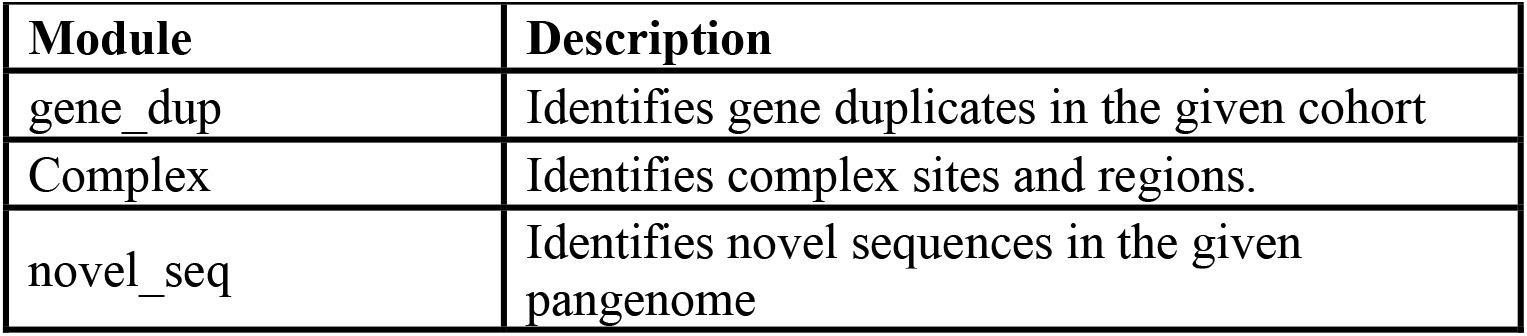
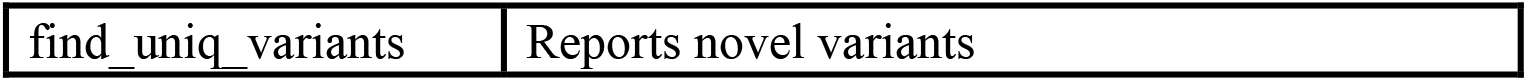
Summary of modules available in PanScan.

**Figure 1.**
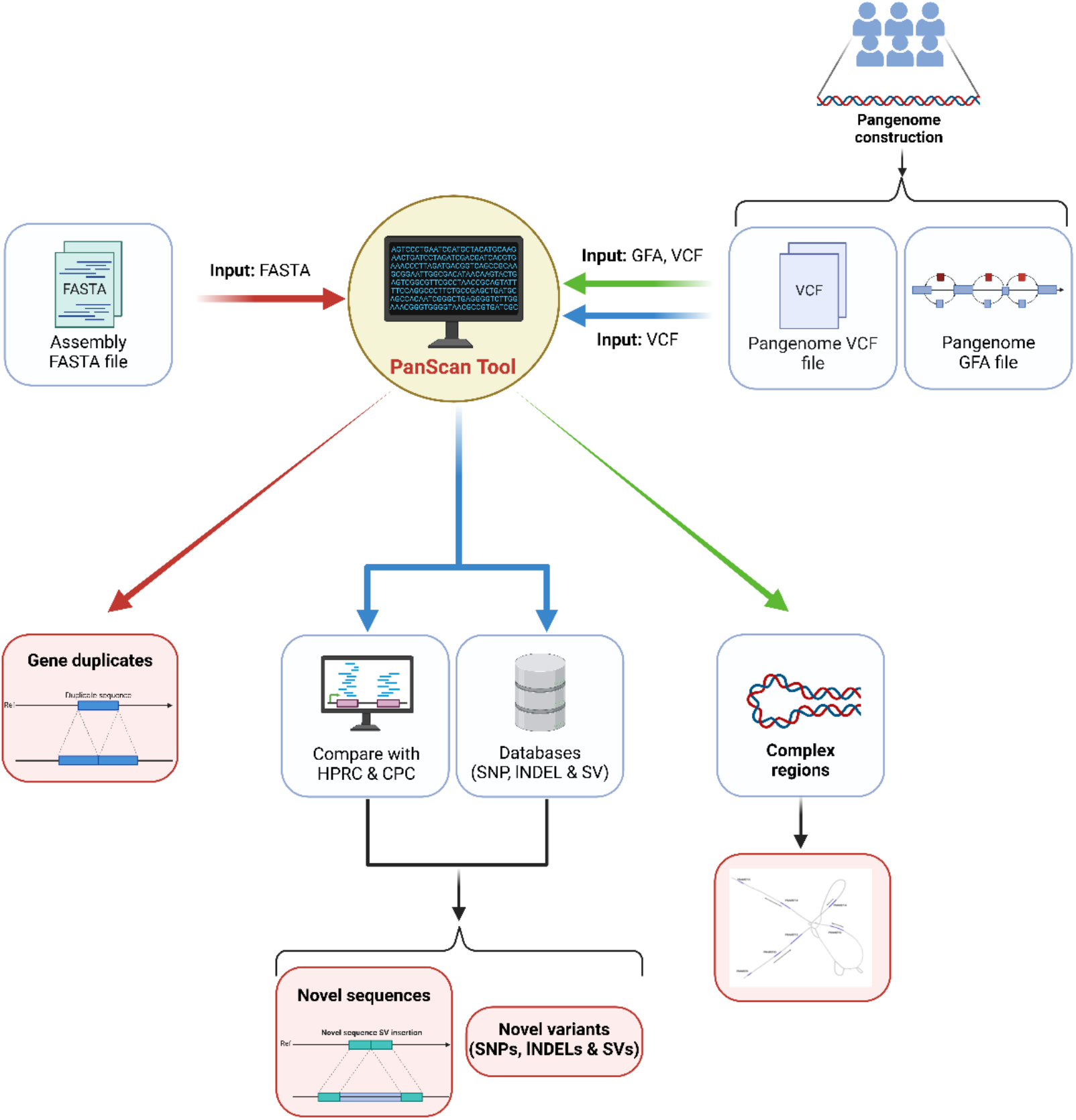
Schematic overview of the PanScan tool and tertiary analysis modules. The diagram illustrates the key input types (FASTA, VCF, and GFA) used for pangenome analysis and their corresponding analytical pathways. Each colored arrow represents a distinct analysis module: red for gene duplication detection, blue for novel sequence & variant identification, and green for structural variant comparison, and complex region analysis. PanScan processes these inputs to compare with published reference datasets (eg: HPRC, CPC), identify novel variants (SNPs, INDELs, and SVs), and analyze complex genomic regions, providing a comprehensive framework for pangenome interpretation.

#### 2.2.1. Gene Duplication

To identify gene duplications in the assembly of interest compared with other reference assemblies, the initial step involves generating a gene duplication matrix for both the query and reference assemblies, which is critical in shaping haplotype architecture. This is particularly relevant for telomere-to-telomere (T2T) assemblies, which address the gaps and limitations present in the GRCh38 reference genome. The generation of the gene duplication matrix is achieved by aligning the assemblies to the GENCODE v38 reference (22) using Liftoff (v1.6.3) (23). Liftoff facilitates the alignment of gene sequences from GENCODE v38 to the target assembly haplotype, optimizing the mapping process to maintain gene structure while maximizing sequencing identity. The following PanScan command is used to generate the gene duplication matrix.

*“panscan make_dup_mtx --gencode_gff3 gencode.v38.annotation.gff3 --fofn assembly_path.txt --ref_fa GRCh38_no_alt_analysis_set.fasta --threads 4”*

The copy number variations (CNVs) for each gene in the target haplotype were then quantitatively assessed by comparing the number of gene copies against the reference (GENCODE v38). PanScan generates the gene duplication matrix in comma-separated values (csv) file format, with the comparison results provided as tab-delimited text files and accompanying images. The workflow for the gene duplication detection module is illustrated in Figure 2A. The PanScan command for analysing gene duplication is provided below:

**Figure 2.**
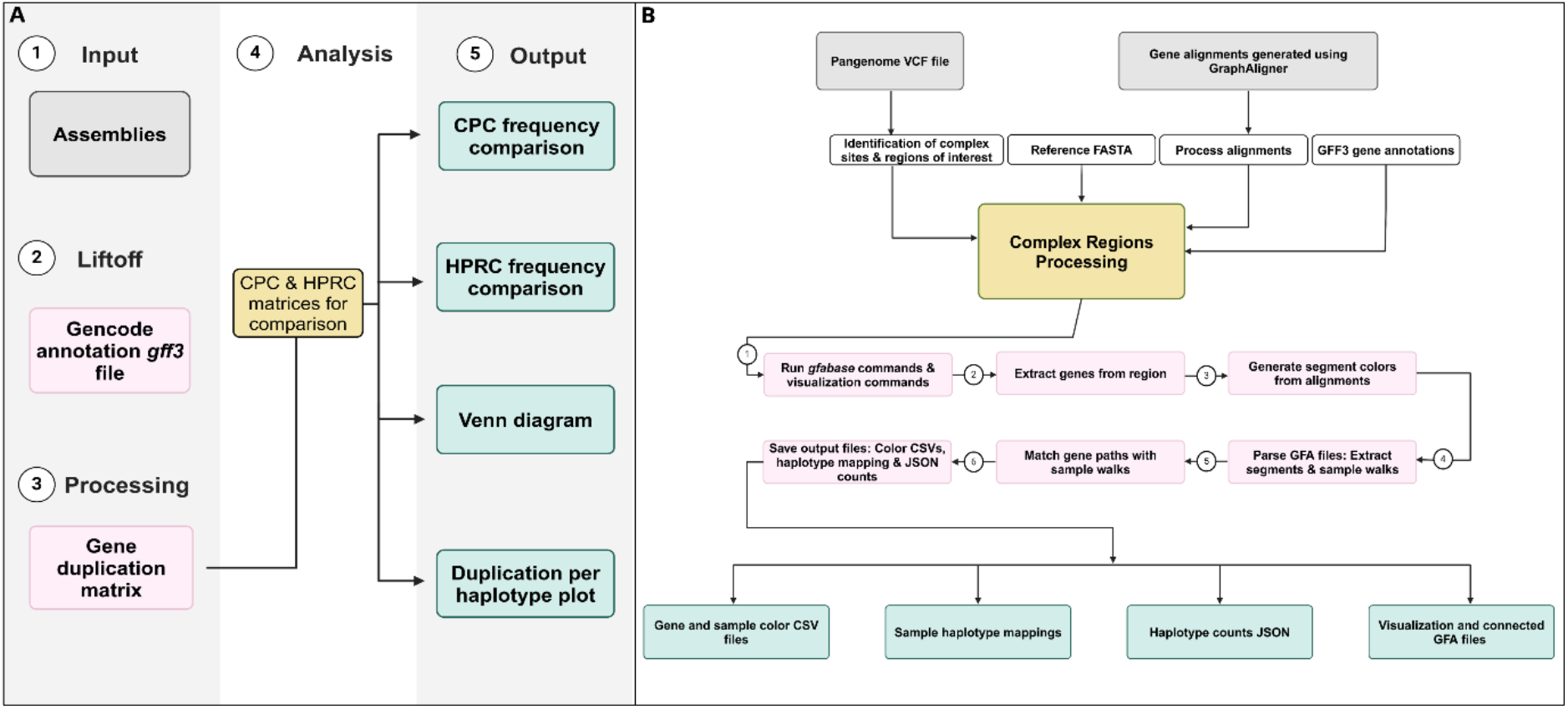
Schematics of workflows for detecting gene duplications and complex regions. (**A**) Workflow for the gene duplication detection module. The figure outlines a step-by-step process, beginning with the input assemblies, followed by gene duplication annotation using Liftoff, processing into a gene duplication matrix, and subsequent comparative analysis with CPC and HPRC reference matrices. The final outputs include frequency comparisons, a Venn diagram, and a duplication per haplotype plot, providing a comprehensive overview of gene duplication events. (**B)** Workflow for identifying complex regions within the studied cohort. The diagram illustrates the stepwise process, starting with the input of pangenome VCF files and gene alignments, followed by the identification and processing of complex regions. The key steps involve extracting gene information from the region, generating segment colors, parsing sample walks, and performing haplotype mapping. The final outputs, including color-coded CSV files, haplotype mappings, JSON counts, and visualization-ready GFA files, aid in the interpretation of complex structural variants (SVs). A two-color scheme is used to differentiate between the input and output stages of the module.

*“panscan gene_dup -i gene-dup-matrix.csv --r1 Reference1-matrix.csv --r2 Reference2-matrix.csv”*

#### 2.2.2. Detection and visualization of complex regions in the pangenome

In the context of pangenome graphs, complex sites are defined as loci harboring at least one SV ≥ 10 kb (default) and at least five independent haplotypes (12). PanScan’s “*complex”* module identifies these sites by analyzing snarls within pangenome VCF files. Unique SVs from the input pangenome GAF file are compared to complex sites. Complex sites with multiple large SVs are deemed “complex structural variation (CSV)” regions if, within a 100kb window, there are at least two multi-allelic complex SV sites, each containing ≥ 10 kb SVs. Gene sequences are mapped to the pangenome graph using GraphAligner (24) with the argument *‘--multimap-score 0.1’*. In cases where multiple genes are mapped to the same locus (in case of isoforms), only the gene with the highest mapping accuracy is retained. To determine whether a haplotype contained a gene, the traversal path is traced through the pangenome graph to see if it contains an intersecting gene region. Variation among individual haplotype walks within the complex region are also visualized using color-coded arrow lines that indicate directionality. The tool generates images for all genes and, additionally, produces plots specifically for the protein-coding genes within the complex region. The workflow for identifying complex regions is illustrated in Figure 2B. The PanScan command for identifying complex regions is:

*“panscan complex --ref_fasta chm13v2.0.fa --gaf_file chm13_mapped_genes.gaf -- sep_pattern ‘#0#’ --gff3 chm13v2.0_RefSeq_Liftoff_v5.1.gff3 -a 5 -n 1 -s 10000 -- regions -l 100000 --sites 1 --sv 1 --ref_name CHM13 --plot_complex panscan.vcf panscan.gfab”*

#### 2.2.3. Identification and visualization of population-specific variants

To streamline and automate multiple layers of variant filtering, we have developed a module that facilitates the detection of novel variants through comparison with existing databases. In PanScan, the *“find_uniq_variants”* module identifies novel SNPs, INDELs, and SVs in the input human pangenome VCF file by comparing them against existing reference (GRCh38) databases (Supplementary Table S1). The process begins with extracting SNPs, INDELs, and SVs greater than 50 bp from the input VCF file for comparison against the databases. In the current version of PanScan, SNPs are compared against dbSNP (25), gnomAD (26), 1000 Genomes (27), and Greater Middle East (GME) (28), while INDELs are compared with gnomAD, 1000 Genomes, and GME. For novel SV identification, PanScan compares input SVs with those reported in the 1000 Genomes and database of genomic variants (DGV) (29) databases. An SV is considered novel if it is not present in the reference databases or if its reciprocal overlap with reference SVs is less than 80% (the default threshold, which can be adjusted using the ‘*--ol’* flag). The workflow for detecting novel variants in a human pangenome VCF file, through a systematic comparison with existing genomic databases, is shown in Figure 3A. The PanScan command for finding novel variants from the given VCF file is:

**Figure 3.**
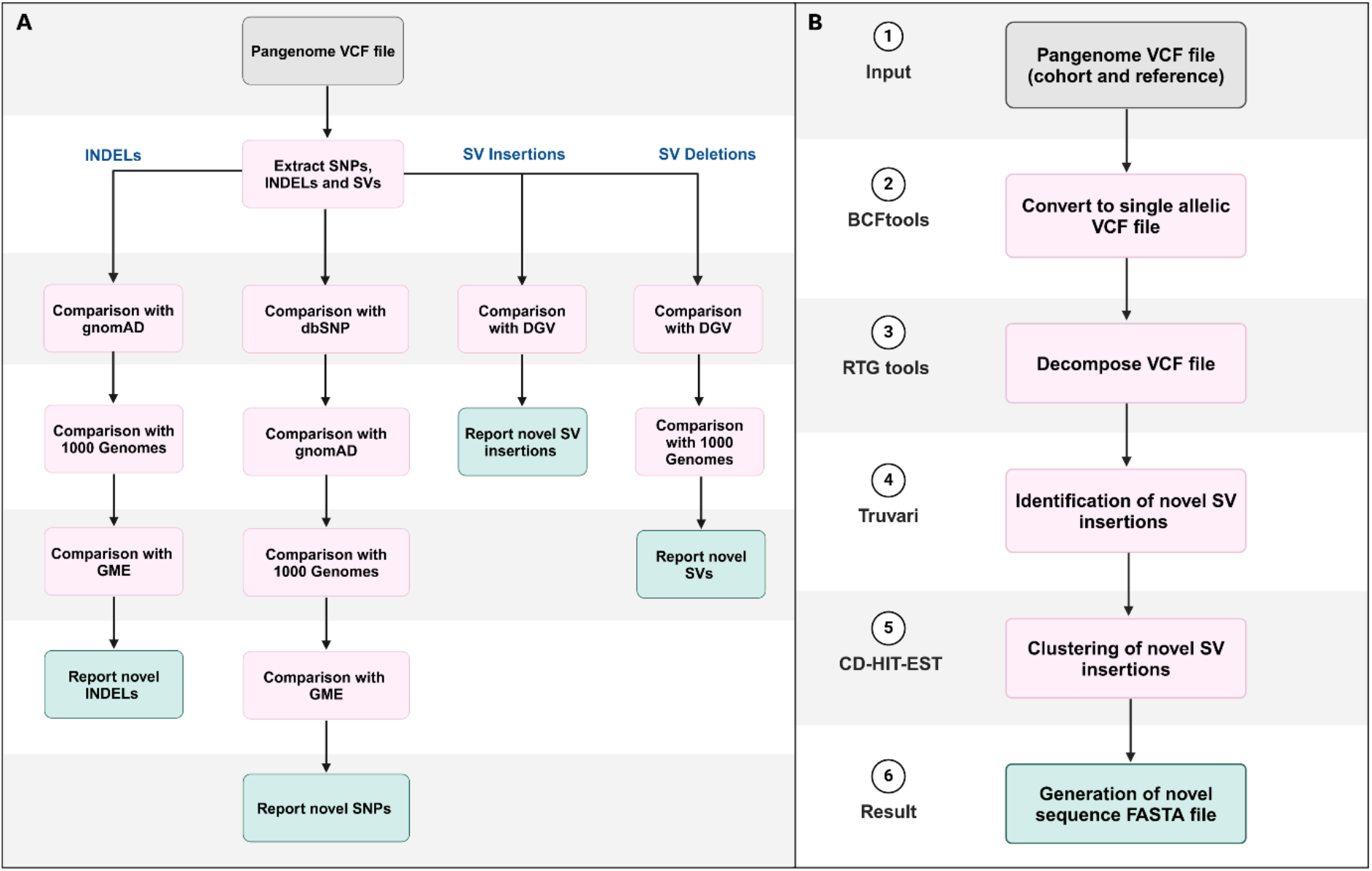
Detailed workflow diagrams representing the process of novel variant discovery and sequence identification across the cohort. **(A)** Workflow for detecting novel variants in a human pangenome VCF file through systematic comparison with existing genomic databases. The process begins with extracting SNPs, INDELs, and SVs from pangenome VCF file, followed by sequential filtering against databases such as gnomAD, dbSNP, 1000 Genomes, GME, and DGV. Novel variants are identified after eliminating known variants, reporting novel SNPs, INDELs, and SVs (insertions and deletions). **(B)** The workflow diagram outlines the steps involved in identifying novel sequences within the studied cohort. The process begins with reading the input pangenome VCF files (both cohort and reference), followed by conversion to single-allelic variants using BCFtools. This is followed by decomposition of complex variants using RTG tools and the identification of unique SV insertions with Truvari. These SV insertions are then clustered using cd-hit-est to generate the final novel sequence FASTA file. Each step in the figure is annotated with the specific tools employed to ensure precise detection and classification of novel sequences.

*“panscan find_uniq_variants -i input.vcf -t SNP --db ALL --overlap 80 –db-path /path/to/databases --op outputdir”*

#### 2.2.4. Identification of novel sequences

Individual haplotypes tend to accumulate new sequence insertions, some of which may be specific to certain populations or individuals. The PanScan *“novel_seq”* module identifies novel sequences within a given pangenome by performing a comparison with a reference pangenome. The workflow begins with the preprocessing of the input and reference VCF files, in which multi-allelic variants are converted into single-allelic variants using the *bcftools norm* command (30). Following this, complex variants are decomposed into SNPs, INDELs (≤50 bp), and SVs (>50 bp) using the *vcf decompose* command from RTG Tools (31). Next, unique SVs in the input pangenome VCF file are identified by comparing them against reference pangenome SVs using the *truvari bench* command from Truvari (32). An SV is classified as unique if its reciprocal overlap with reference SVs is below 80%, ensuring that multiple overlapping SVs are consolidated into a single representative variant. Finally, the identified unique SV insertions are clustered using CD-HIT-EST (version 4.8.1) (33, 34) with a default sequence identity threshold of 0.9, and the resulting clusters are reported as novel sequences (Supplementary Figure S1). An overview of the novel sequence identification workflow is provided in Figure 3B. The PanScan command for identifying novel sequences in the provided pangenome VCF file is:

*“panscan novel_seq -i input.vcf -r reference.vcf -t 4 --dpi 600 --op outputdir”*

### 2.3. Software dependencies

PanScan comprises multiple heuristic scripts, each executing a combination of custom Python and Perl scripts along with commands from third-party software. These include Liftoff (23), GraphAligner (24), GFABase (https://github.com/mlin/gfabase), Bandage (35), BCFtools (30), RTG Tools (31), Truvari (32), BGZIP (36), Tabix (37), and CD-HIT-EST (33, 34), as well as widely used Python libraries such as NumPy, Pandas, and Pickle. Additionally, a Java class file (reciprocal_overlap.class) was implemented in the *“find_uniq_variants”* module to assess the reciprocal overlap of SVs. The Rideogram (38) package in R was used to generate ideogram visualizations. A comprehensive list of third-party tools, along with their references, is provided in Supplementary Table S2. PanScan is openly accessible on GitHub (https://github.com/CATG-Github/panscan) and includes comprehensive documentation covering installation, configuration, and usage.

### 2.4. Assessment of compute time, memory usage, and disk space

The UNIX time command was utilized to evaluate program performance, where ‘Elapsed (wall clock) time’ measures total execution duration, and ‘Maximum resident set size (kbytes)’ quantifies peak memory utilization. To estimate the average runtime of each module in PanScan using the GIAB test dataset, log files were analysed instead. All analyses were computed on the same server running Rocky Linux 8.10 with Intel(R) Xeon(R) Gold 5220 CPU @ 2.20GHz processors, 64 GB of random-access memory, and 2.2 Petabyte storage space. The compute time and resource usage for each PanScan module are provided in Supplementary Table S3.

## 3. Results

### 3.1. Pangenome graph of GIAB dataset

We constructed a pangenome graph to demonstrate the utility and tertiary analysis capabilities of the PanScan tool by incorporating three trios from the GIAB dataset, (HG002, HG005, and NA19240) which were not included in the HPRC pangenome. The resulting pangenome encompassed a total length of 3,164,974,913 base pairs, comprising 32,199,662 nodes and 43,740,708 edges. To characterize genetic variation in this cohort, we compared the GIAB dataset pangenome graph to the combined CPC-HPRC (Chinese Pangenome Consortium and Human Pangenome Reference Consortium) pangenome (10, 11), a resource encompassing diverse human populations. Further analysis of the applicability of the PanScan tool and its outputs, including those derived from the GIAB dataset is presented in the following sections.

### 3.2. Gene duplications in the GIAB set assemblies

PanScan uses Liftoff v1.6.3 tool (23) to identify the gene duplication events within the diploid assemblies of the three GIAB trios and GENCODE v38 (https://www.gencodegenes.org/human/) was used for the gene annotation. This approach facilitated the identification of extensive gene duplications, thereby uncovering novel and diverse genomic features.

A comparative assessment with respect to HPRC and CPC assemblies revealed duplicated genes unique to the GIAB assemblies that were absent from the HPRC assemblies and a further 1,351 duplicated genes that were absent from both the HPRC and CPC assemblies (Figure 4A). In addition, 12 duplicated genes from the GIAB assemblies were also observed in the HPRC and CPC assemblies. Among these overlapping genes, some genes were present at a greater frequency (≥5%) in the GIAB dataset assemblies than in the HPRC (Figure 4B), such as the H3-2 gene. However, the GIAB assemblies did not have any genes that were duplicated more frequently than CPC, if there would have been, the tool would have produced a comparative plot. Recurrent gene duplications specific to GIAB assemblies further highlight their unique genetic landscape, with 120 and 64 duplicated genes not found in the HPRC and CPC assemblies, respectively (Figure 4A). In addition, 13 duplicated genes from the GIAB assemblies were also observed in the HPRC and CPC assemblies. The gene duplication matrix for the GIAB dataset after reference correction is provided in Supplementary Table 4A. Supplementary Tables 4B and 4C present the frequency comparisons of common gene duplicates between the GIAB dataset and the HPRC and CPC cohorts.

**Figure 4.**
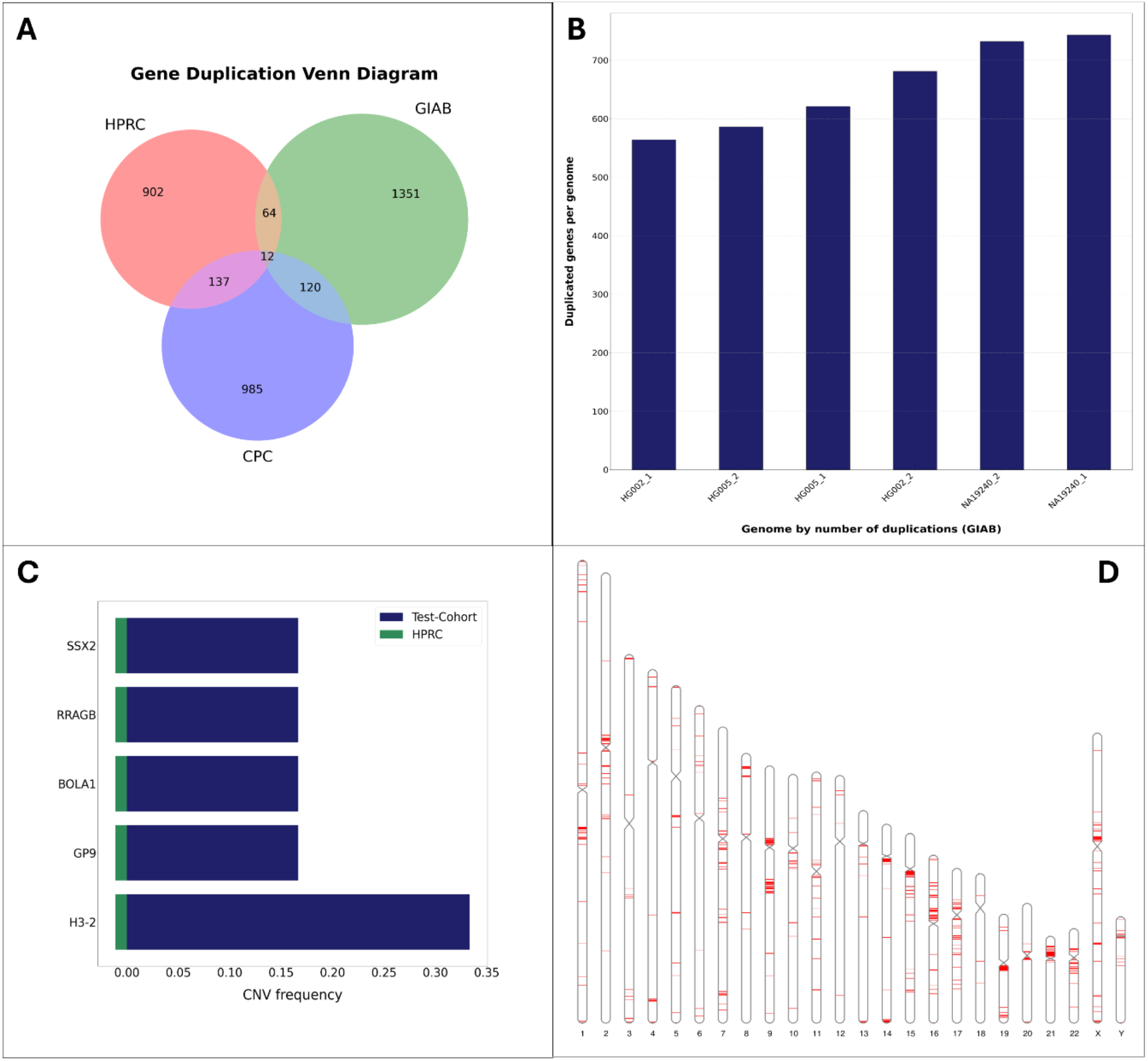
PanScan output from the “*gene_dup”* module. (**A**) Venn diagram illustrating the overlap of gene duplications among GIAB, HPRC and CPC cohorts. (**B**) Bar plot displaying the most frequently duplicated genes in the GIAB cohort compared to HPRC. (**C**) Bar plot showing the number of duplicated genes per assembly in the GIAB dataset. (**D**) Ideogram representing the chromosomal locations of gene duplications, highlighting their distribution across the genome.

### 3.3. Complex SV regions in the GIAB pangenome graph

We used a pangenome graph to visualize and analyze CSV in the GIAB samples. In PanScan, by default, a CSV site is defined as an SV site that contains at least one SV of minimum 10 kb in length and a minimum of 5 haplotypes passing through it. Our analysis revealed that most CSVs were approximately 10 kb in length, which was the established lower bound for CSV inclusion in our analysis, as mentioned in our previous study. Utilizing rigorous criteria, we delineated CSV regions as areas within a 100 kb window that consisted of at least one CSV site and an additional SV. From the dataset, PanScan identified 49 complex regions, within which 43 genes were detected (Supplementary Table 5A & 5B). Figure 5 displays the visualization of complex genomic regions using PanScan. Supplementary Figure S2 shows the paths traversed by the sample haplotypes and the reference genomes (CHM13 and GRCh38) within a representative complex region.

**Figure 5.**
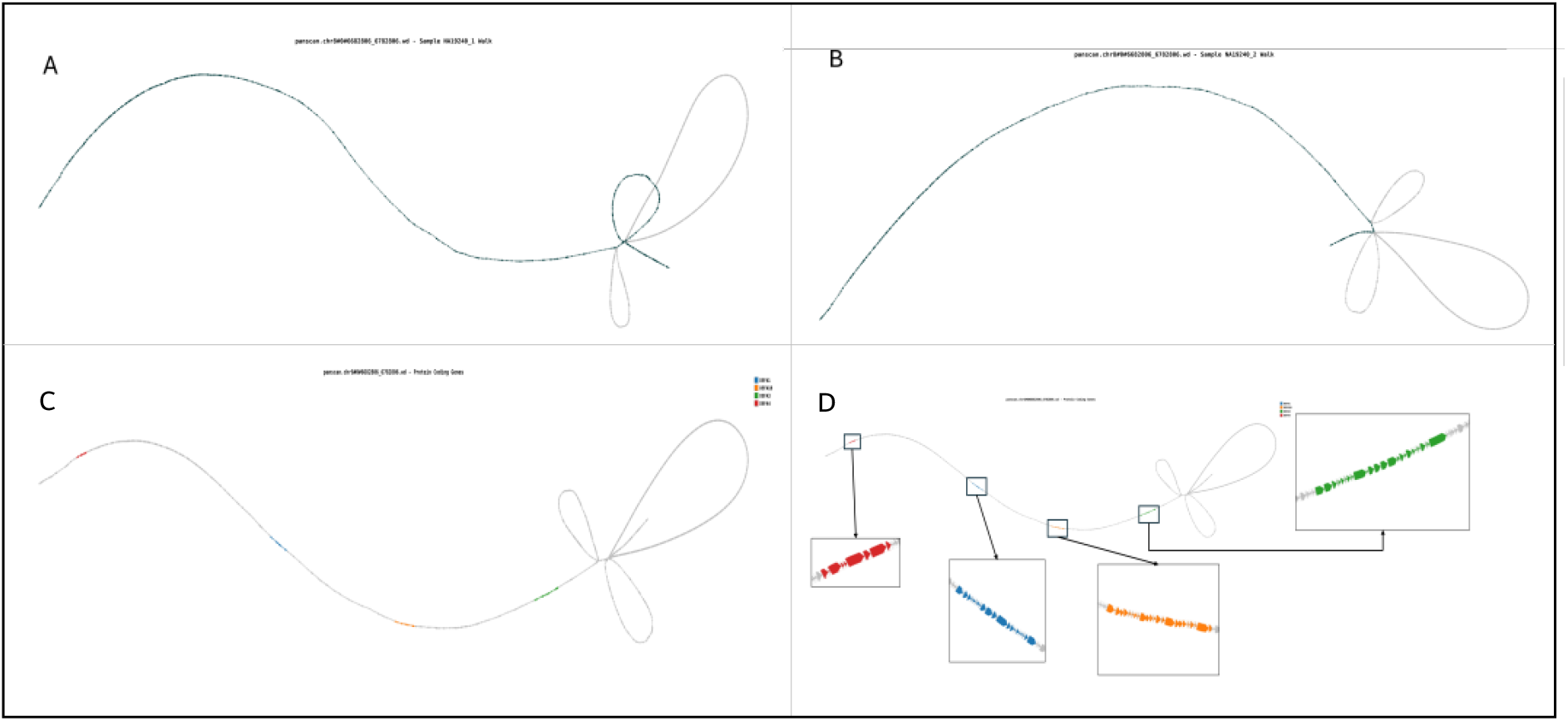
Visualization of the complex regions using PanScan. The subgraphs of the pangenome containing the CSV regions are loaded into Bandage along with the respective color code (e.g. ‘#ff00007’) for the distinct protein coding genes (if any) and the traversal path is traced based on the color gradient from red to blue.

### 3.4. Discovery of Novel Variants Using PanScan

To evaluate the performance of PanScan, we applied it to a pangenome VCF file constructed from three assemblies and conducted a stepwise comparison with established variant databases, including dbSNP, gnomAD, 1000 Genomes, and GME for SNPs and INDELs, as well as DGV and 1000 Genomes for SVs (Supplementary Tables S6A, S6B & S6C). This approach facilitated the systematic exclusion of known variants, ensuring that only novel variants were retained.

For SNPs, a total of 7,989,787 variants were initially analyzed. Using PanScan, the dataset was first compared to dbSNP, identifying 204,973 common SNPs which were subsequently removed. The remaining 7,784,814 SNPs were then filtered against gnomAD, resulting in the elimination of 985 additional common SNPs. A comparison with 1000 Genomes led to the removal of 934 further SNPs. Finally, the remaining 7,782,895 SNPs were compared to GME, resulting in the exclusion of 34 common SNPs. Following this systematic elimination process, 7,782,861 SNPs were classified as novel, as they were not cataloged in any of the four reference databases. This stepwise approach highlights the limited overlap between known genomic databases and the significant number of uncharacterized SNPs that PanScan successfully identified.

A similar strategy was employed for INDELs. Of the 2,751,986 INDELs detected, PanScan identified 92 variants common to gnomAD, which were subsequently excluded. Comparisons with 1000 Genomes and GME revealed no additional overlaps, resulting in a final count of 2,751,894 novel INDELs. The minimal overlap with known databases suggests that current genomic reference datasets may underrepresent small insertions and deletions.

PanScan identified a similarly high proportion of novel events for SVs. Among 33,846 deletions, only 16 were present in DGV, and 2 in 1000 Genomes, resulting in 33,828 novel deletions. Similarly, of the 42,373 insertions identified, only 19 were previously cataloged, leaving 42,354 novel insertions. These results suggest that the current SV databases account for only a small fraction of large genomic alterations, highlighting the need for more extensive and comprehensive SV datasets.

The results obtained through PanScan highlight the gaps in the current genomic reference panels. While dbSNP showed the highest overlap with the input SNPs, the majority of identified variants were not previously reported. gnomAD, 1000 Genomes, and GME provided only minimal additional filtering, further underscoring the novelty of the discovered variants. The identification of 7,782,861 novel SNPs, 2,751,894 novel INDELs, and over 76,000 novel SVs demonstrates that a pangenome-based approach facilitates the discovery of previously uncharacterized genetic diversity on a large scale.

### 3.5. Novel Sequences

Using the “novel_seq” module of PanScan, we identified novel sequences in the GIAB datasets by comparing SV insertions against the CPC and HPRC pangenome. The analysis produced a FASTA file containing the novel sequences, along with a text file ranking the samples in ascending order based on the number of novel sequences identified. In total, PanScan reported 17,218 novel sequences comprising 18,806,147 base pairs from the GIAB dataset. The novel sequence fasta file is available in Zenodo (https://doi.org/10.5281/zenodo.15301483). The distribution included 8,624 sequences from sample NA19240, 5,863 from HG005, and 5,667 from HG002. The GRCh38 sample, which served as one of the reference genomes during pangenome construction, was excluded from this analysis using the “--exclude GRCh38” option in the “novel_seq” command. The output also included comprehensive genomic annotations for each novel sequence, such as chromosome coordinates, variant positions, SV insertion lengths, and sample-wise presence. Furthermore, the analysis indicated whether each novel sequence was shared across multiple samples. An ideogram was also generated to visualize the chromosomal distribution of these sequences, providing a clear representation of their genomic localization.

## Discussion

PanScan is a robust and user-friendly tertiary analysis tool developed for graph-based pangenome interpretation. With pangenomes representing a recent advancement in genomic analysis, PanScan provides an accessible and efficient framework for both clinical and research applications. Graph-based pangenome analysis has significantly improved our ability to identify complex genomic regions, novel sequences, and population-specific variations, providing a more comprehensive view of human genetic diversity. We demonstrate how PanScan, a newly developed computational tool, effectively and efficiently captures genetic variation from regions not well represented by traditional linear reference genomes, making it a scalable and powerful tool for pangenome interpretation. By analysing the GIAB dataset against the CPC-HPRC pangenome reference, we identified numerous novel sequences, population-specific SVs, gene duplication events, and highly complex genomic regions, thus showcasing the usability of the tool. These findings emphasize the importance of integrating graph-based pangenomes into standard genomic analyses to enhance the resolution of SVs and haplotype complexity.

In this study, we constructed a pangenome graph using Minigraph-Cactus, incorporating six haplotypes from the GIAB dataset, CHM13 as the scaffold, and the GRCh38 reference genome. The resulting structure encompassed 3.16 billion based pairs with over 32 million nodes and 43 million edges, making the graph highly dense and interconnected for comparative genomic analysis (11, 21). Unlike linear reference genomes, which often suffer from reference bias and fail to fully capture genetic diversity, pangenome approaches provide a more accurate and comprehensive representation of genomic variation. By enabling the detection of novel sequences, gene duplications, and complex SVs, PanScan facilitates deeper insights into genomic architecture, reinforcing the need for graph-based pangenomes in advancing human genetic research.

## Code Availability

The source code for PanScan is available on the GitHub repository at: https://github.com/CATG-Github/panscan. A comprehensive manual for all PanScan functions, along with vignettes demonstrating their functionality, and links to use cases and benchmarking code, can also be found on the PanScan GitHub page. The pre-processed reference VCF files used for benchmark are uploaded to Zenodo (https://doi.org/10.5281/zenodo.15301483).

## Acknowledgments

We would like to express our gratitude to Prof. Alawi Alsheikh-Ali and Prof. Stefan S. Du Plessis for their support in securing the grant for this study. We are also grateful to the Center for Applied and Translational Genomics (CATG) at Mohammed Bin Rashid University of Medicine and Health Sciences (MBRU), Dubai Health, Dubai, UAE, for providing the computational infrastructure and resources for the development of this tool.

## Author Contribution

MU conceived the name, conceptualized and supervised the study. BB, SH, MAH and MK made equal contributions to this work. BB, SH, MAH, and MK developed the code, designed the methodology, and wrote the original draft. SB created the workflow images. MAH, BB, and SH analysed the data and created the figures and tables. MM, BJ, and NN contributed to the original draft. SS assisted with formatting and referencing. RT and TA conducted the tool testing. HE, MAO, NM, KDHP, AIH, NCS, OA, SSDP, MA, AA, and MU reviewed, edited, and approved the final version of the manuscript.

## Supplementary data

Supplementary Data are available at NAR online.

## References

1. Lander, E.S., Linton, L.M., Birren, B., Nusbaum, C., Zody, M.C., Baldwin, J., Devon, K., Dewar, K., Doyle, M., FitzHugh, W., et al. (2001) Initial sequencing and analysis of the human genome. Nature, 409, 860–921.

2. Abondio, P., Cilli, E. and Luiselli, D. (2023) Human Pangenomics: Promises and Challenges of a Distributed Genomic Reference. Life (Basel), 13.

3. Golicz, A.A., Bayer, P.E., Bhalla, P.L., Batley, J. and Edwards, D. (2020) Pangenomics Comes of Age: From Bacteria to Plant and Animal Applications. Trends Genet, 36, 132–145.

4. Hurgobin, B. and Edwards, D. (2017) SNP Discovery Using a Pangenome: Has the Single Reference Approach Become Obsolete? Biology (Basel), 6.

5. Stenman, A., Yang, M., Paulsson, J.O., Zedenius, J., Paulsson, K. and Juhlin, C.C. (2021) Pan-Genomic Sequencing Reveals Actionable CDKN2A/2B Deletions and Kataegis in Anaplastic Thyroid Carcinoma. Cancers (Basel), 13.

6. Yu, Y., Zhang, Z., Dong, X., Yang, R., Duan, Z., Xiang, Z., Li, J., Li, G., Yan, F., Xue, H., et al. (2022) Pangenomic analysis of Chinese gastric cancer. Nat Commun, 13, 5412.

7. Tay Fernandez, C.G., Nestor, B.J., Danilevicz, M.F., Marsh, J.I., Petereit, J., Bayer, P.E., Batley, J. and Edwards, D. (2022) Expanding Gene-Editing Potential in Crop Improvement with Pangenomes. Int J Mol Sci, 23.

8. Gao, Y., Guitton-Sert, L., Dessapt, J., Coulombe, Y., Rodrigue, A., Milano, L., Blondeau, A., Larsen, N.B., Duxin, J.P., Hussein, S., et al. (2023) A CRISPR-Cas9 screen identifies EXO1 as a formaldehyde resistance gene. Nat Commun, 14, 381.

9. Nassir, N. A., Almarri, M., Akter, H., Hassan Khansaheb, H., Uddin, K.M.F., Abou Tayoun, A., Du Plessis, S.S., Haber, M., Alsheikh-Ali, A. and Uddin, M. (2025) Advancing clinical genomics with Middle Eastern and South Asian pangenomes. Nat Med, 31.

10. Liao, W.-W., Asri, M., Ebler, J., Doerr, D., Haukness, M., Hickey, G., Lu, S., Lucas, J.K., Monlong, J., Abel, H.J., et al. (2023) A draft human pangenome reference. Nature, 617, 312–324.

11. Gao, Y., Yang, X., Chen, H., Tan, X., Yang, Z., Deng, L., Wang, B., Kong, S., Li, S., Cui, Y., et al. (2023) A pangenome reference of 36 Chinese populations. Nature, 619, 112–121.

12. Nassir, N., Almarri, M.A., Kumail, M., Mohamed, N., Balan, B., Hanif, S., AlObathani, M., Jamalalail, B., Elsokary, H., Kondaramage, D., et al. (2024) A draft Arab pangenome reference. bioRxiv, 10.1101/2024.07.09.602638.

13. Littlefield, C., Lazaro-Guevara, J.M., Stucki, D., Lansford, M., Pezzolesi, M.H., Taylor, E.J., Wolfgramm, E.-M.C., Taloa, J., Lao, K., Dumaguit, C.D.C., et al. (2024) A Draft Pacific Ancestry Pangenome Reference. bioRxiv, 10.1101/2024.08.07.606392.

14. Sheikhizadeh, S., Schranz, M.E., Akdel, M., de Ridder, D. and Smit, S. (2016) PanTools: representation, storage and exploration of pan-genomic data. Bioinformatics, 32, i487–i493.

15. Ding, W., Baumdicker, F. and Neher, R.A. (2018) panX: pan-genome analysis and exploration. Nucleic Acids Res, 46, e5.

16. Cannon, S.B., Lee, H.-O., Weeks, N.T. and Berendzen, J. (2024) Pandagma: a tool for identifying pan-gene sets and gene families at desired evolutionary depths and accommodating whole-genome duplications. Bioinformatics, 40.

17. Noll, N., Molari, M., Shaw, L.P. and Neher, R.A. (2023) PanGraph: scalable bacterial pan-genome graph construction. Microb Genom, 9.

18. Li, J., Yang, A., Carneiro, B.A., Gamsiz Uzun, E.D., Massingham, L. and Uzun, A. (2024) Variant graph craft (VGC): a comprehensive tool for analyzing genetic variation and identifying disease-causing variants. BMC Bioinformatics, 25, 288.

19. Goel, M., Sun, H., Jiao, W.-B. and Schneeberger, K. (2019) SyRI: finding genomic rearrangements and local sequence differences from whole-genome assemblies. Genome Biol, 20, 277.

20. Marçais, G., Delcher, A.L., Phillippy, A.M., Coston, R., Salzberg, S.L. and Zimin, A. (2018) MUMmer4: A fast and versatile genome alignment system. PLoS Comput Biol, 14, e1005944.

21. Hickey, G., Monlong, J., Ebler, J., Novak, A.M., Eizenga, J.M., Gao, Y., Abel, H.J., Antonacci-Fulton, L.L., Asri, M., Baid, G., et al. (2023) Pangenome graph construction from genome alignments with Minigraph-Cactus. Nature Biotechnology 2023 42:4, 42, 663–673.

22. Mudge, J.M., Carbonell-Sala, S., Diekhans, M., Martinez, J.G., Hunt, T., Jungreis, I., Loveland, J.E., Arnan, C., Barnes, I., Bennett, R., et al. (2025) GENCODE 2025: reference gene annotation for human and mouse. Nucleic Acids Res, 53, D966–D975.

23. Shumate, A. and Salzberg, S.L. (2021) Liftoff: accurate mapping of gene annotations. Bioinformatics, 37, 1639–1643.

24. Rautiainen, M. and Marschall, T. (2020) GraphAligner: rapid and versatile sequence-to-graph alignment. Genome Biol, 21, 253.

25. Phan, L., Zhang, H., Wang, Q., Villamarin, R., Hefferon, T., Ramanathan, A. and Kattman, B. (2025) The evolution of dbSNP: 25 years of impact in genomic research. Nucleic Acids Res, 53, D925–D931.

26. Karczewski, K.J., Francioli, L.C., Tiao, G., Cummings, B.B., Alföldi, J., Wang, Q., Collins, R.L., Laricchia, K.M., Ganna, A., Birnbaum, D.P., et al. (2020) The mutational constraint spectrum quantified from variation in 141,456 humans. Nature, 581, 434–443.

27. Highnam, G., Franck, C., Martin, A., Stephens, C., Puthige, A. and Mittelman, D. (2013) Accurate human microsatellite genotypes from high-throughput resequencing data using informed error profiles. Nucleic Acids Res, 41.

28. Scott, E.M., Halees, A., Itan, Y., Spencer, E.G., He, Y., Azab, M.A., Gabriel, S.B., Belkadi, A., Boisson, B., Abel, L., et al. (2016) Characterization of Greater Middle Eastern genetic variation for enhanced disease gene discovery. Nat Genet, 48, 1071–1079.

29. MacDonald, J.R., Ziman, R., Yuen, R.K.C., Feuk, L. and Scherer, S.W. (2014) The Database of Genomic Variants: a curated collection of structural variation in the human genome. Nucleic Acids Res, 42.

30. Danecek, P., Bonfield, J.K., Liddle, J., Marshall, J., Ohan, V., Pollard, M.O., Whitwham, A., Keane, T., McCarthy, S.A. and Davies, R.M. (2021) Twelve years of SAMtools and BCFtools. Gigascience, 10.

31. Cleary, J.G., Braithwaite, R., Gaastra, K., Hilbush, B.S., Inglis, S., Irvine, S.A., Jackson, A., Littin, R., Rathod, M., Ware, D., et al. (2015) Comparing Variant Call Files for Performance Benchmarking of Next-Generation Sequencing Variant Calling Pipelines. bioRxiv, 10.1101/023754.

32. English, A.C., Menon, V.K., Gibbs, R.A., Metcalf, G.A. and Sedlazeck, F.J. (2022) Truvari: refined structural variant comparison preserves allelic diversity. Genome Biol, 23.

33. Li, W. and Godzik, A. (2006) Cd-hit: a fast program for clustering and comparing large sets of protein or nucleotide sequences. Bioinformatics, 22, 1658–1659.

34. Fu, L., Niu, B., Zhu, Z., Wu, S. and Li, W. (2012) CD-HIT: accelerated for clustering the next-generation sequencing data. Bioinformatics, 28, 3150–3152.

35. Wick, R.R., Schultz, M.B., Zobel, J. and Holt, K.E. (2015) Bandage: interactive visualization of de novo genome assemblies. Bioinformatics, 31, 3350–3352.

36. Bonfield, J.K., Marshall, J., Danecek, P., Li, H., Ohan, V., Whitwham, A. and Keane, T. (2021) HTSlib: C library for reading/writing high-throughput sequencing data. Gigascience, 10, giab007.

37. Li, H. (2011) Tabix: fast retrieval of sequence features from generic TAB-delimited files. Bioinformatics, 27, 718–719.

38. Hao, Z., Lv, D., Ge, Y., Shi, J., Weijers, D., Yu, G. and Chen, J. (2020) RIdeogram: Drawing SVG graphics to visualize and map genome-wide data on the idiograms. PeerJ Comput Sci, 6, 1–11.

